# Switching On Static Gene Regulatory Networks to Compute Cellular Decisions

**DOI:** 10.1101/2020.05.29.122200

**Authors:** Clara E. Pavillet, Dimitrios Voukantsis, Francesca M. Buffa

## Abstract

**Motivation:** Gene networks are complex sets of regulators and interactions that govern cellular processes. Their perturbations can disrupt regular biological functions, translating into a change in cell behaviour and ability to respond to internal and external cues. Computational models of these networks can boost translation of our scientific knowledge into medical applications by predicting how cells will behave in health and disease, or respond to stimuli such as a drug treatment. The development of such models requires effective ways to read, manipulate and analyse the increasing amount of existing, and newly deposited gene network data.

**Results:** We developed BioSWITCH, a command-line program using the BioPAX standardised language to “switch on” static regulatory networks so that they can be executed in GINML to predict cellular behaviour. Using a previously published haematopoiesis gene network, we show that BioSWITCH successfully and faithfully automates the network de-coding and re-coding into an executable logical network. BioSWITCH also supports the integration of a BioPAX model into an existing GINML graph.

**Availability:** Source code available at https://github.com/CBigOxf/BioSWITCH.

## 1 Introduction

Complex inner-cell molecular networks and their dynamical properties underpin cellular behaviour and interactions with the surrounding environment. The ability to model such network has the potential to boost our understanding of health and disease and translate our knowledge into useful clinical tools. We and others have recently illustrated how a computational technique such as Agent-Based Modelling (ABM) lends itself naturally to represent cells, as computational *meta-agents*, in a three-dimensional (3D) environment equipped with an inner network model of genes and molecules (Voukantsis et al. 2019, Letort et al. 2019). As a result, co-occurring intrinsic molecular changes and extrinsic factors influencing cellular behaviour can be modelled within the broader biological context of the environment in which they act.

Achieving a meaningful link between genotype and phenotype via gene networks, however, requires careful curation of deposited and newly acquired biological data. The increased availability of high-throughput technology has boosted our ability to acquire such data, and led to the creation of large data repositories, including pathway repositories such as Reactome (Fabregat et al., 2016), WikiPathways (Kutmon et al., 2016), Pathway Commons (Cerami et al., 2011), and KEGG (Kanehisa et al., 2017). These are assimilated by their aim to provide easy access to biological entities and their corresponding interactions, and play a vital role in the construction of comprehensive models requiring the manual or computational extraction of information from literature and databases. A caveat is the fractionation of biological data into a variety of languages and formats across different data providers. This translates into computational tools supporting the analysis of data written in some formats but not others, a phenomenon that creates redundancy and compatibility issues, highlighting the need for standardisation through the development of a widely adopted format. Although there may never be an Esperanto language, the Biological Pathway Exchange (BioPAX) language (Demir et al., 2010), an OWL-based language represented in the RDF/XML format (Horrocks et al., 2003), represents a community-driven effort to increase uniformity of pathway data. Since its creation it is becoming a widely used format and major databases increasingly support file export to BioPAX. Software like Chisio BioPAX Editor (ChiBE) (Babur et al., 2010) and Cytoscape (Shannon P, 2003), along with parsing tools like Paxtools (Demir et al., 2013) and PaxtoolsR (Luna et al., 2016), enable the visualisation and parsing of these models, respectively.

BioPAX networks however are *static*, with no established method supporting their execution and generation of testable predictions. With the aim to develop BioPAX-compatible executable pathway models, Haydarlou et al. (2016) created a Java framework, BioASF, which partially addresses the above. However, the network simulation rules need to be manually encoded in Java when a new model is developed, and no user interface exists, which limits non-developers’ ability to generate new models or expand them. BioSWITCH was developed to address these limitations, and to automate the translation of BioPAX graphs into executable models containing the necessary logical rules to be simulated in different environments (i.e. turned *ON*). Importantly, the simulation rules are generated directly from the BioPAX functional annotation of the pathways, with no additional coding needed by the user. As we continue to further develop BioSWITCH, we will increasingly be able to tap into the comprehensive collection of biological data in the form of annotated graphs, thereby facilitating the automated incorporation and scaling of executable predictive models.

## 2 Description

### 2.1 Extracting Data from BioPAX Gene Regulatory Networks to Construct Executable Models

BioSWITCH is a command-line application developped in Python. It produces networks with directed edges in the GINML format (Naldi et al., 2009), an extension of the Graph eXchange Language (GXL) (Holt et al., 2006). It takes in a BioPAX Gene Regulatory Network model containing the ontology classes in Figure 1a and extracts the graph ID, the specific ontology terms for the biological entities and their interactions. It then converts these into specific rules which can be used for the simulation of the network. Specifically, once a *BiopaxGinml* object has been initialised, the BioSWITCH function *WriteGINML* automatically generates a logical GINML regulatory graph (cf. Figure 1b).

**Fig. 1.**
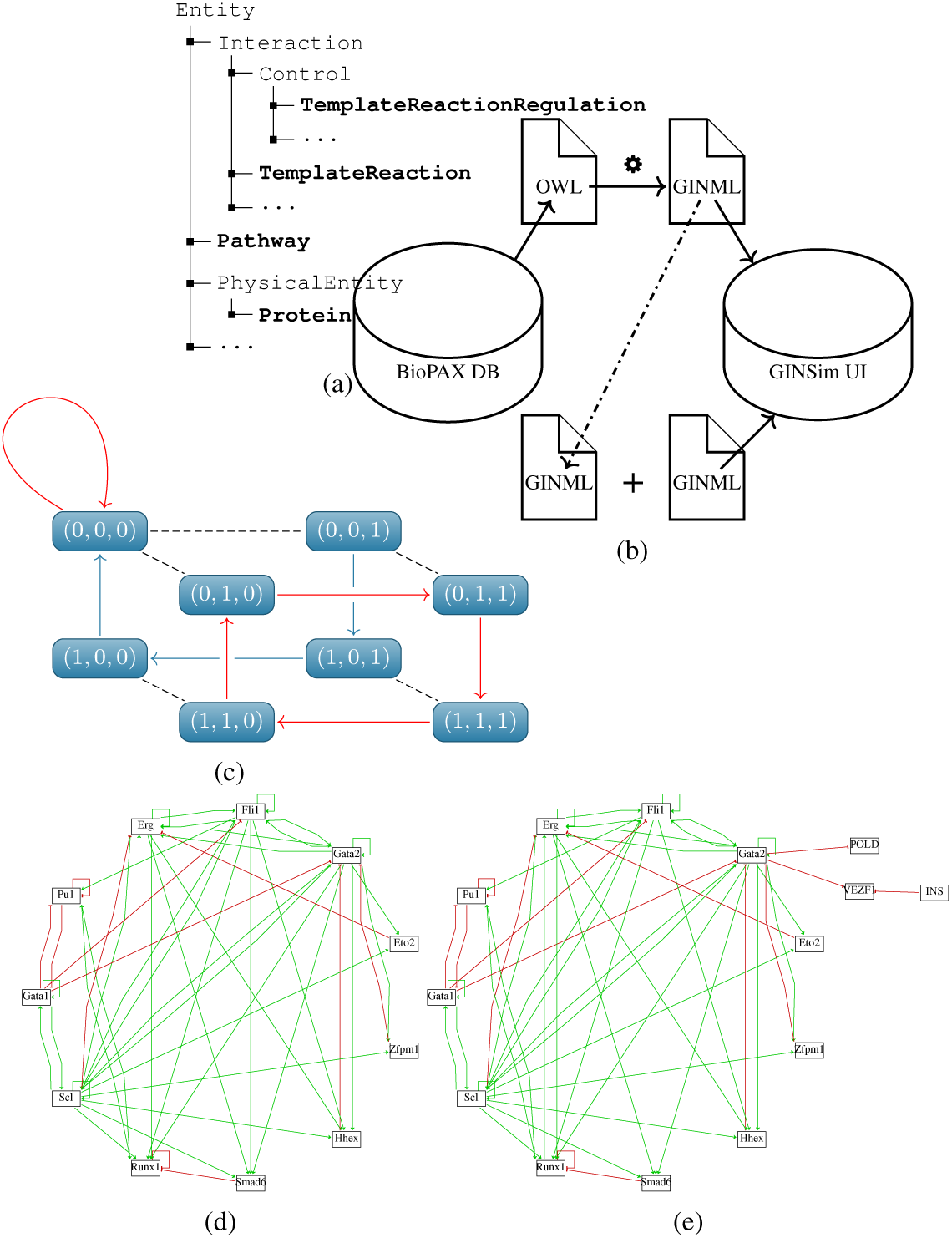
a) Overview of the BioPAX ontology classes currently supported by BioSWITCH. b) BioSWITCH workflow, enabling the conversion of a BioPAX model (OWL) to GINML, with the optionality of directly integrating it into an existing GINML network. c) Schematic diagram representing the state space of a Boolean model composed of 3 nodes. Stable states are shown in red. d) Generated haematopoiesis GINML model by BioSWITCH in GINSim. e) Figure d merged by BioSWITCH with a smaller BioPAX network consisting of three additional nodes and one overlapping node.

### 2.2 Simulating Gene Regulatory Networks

The generated GINML file encodes both the information required to create a visual representation and the logical formalism for the network to be executed. A handful of tools exist to execute biological networks, the Gene Interaction Network Simulation (GINSim) environment, a freeware Java application (Naldi et al., 2009), offers a user-friendly graph editor for the manual creation and/or editing of models, and provides a database to share the models for reuse by the scientific community.

A network, or graph, *G* = (*N, E*), is formally defined by a set of objects, or nodes, (*N*) connected together by links, or edges (*E*). In GINSim, simple geometric vector shapes define the nodes (ie. genes, proteins), whilst line segments drawn between nodes define the interaction types (ie. activating → or inhibitory –|). Beyond this visual structure, logical parameters can be defined governing the rules for activation of individual nodes in the network, using for example, but not limited to, the Boolean expressions: & = AND, ! = NOT, | = OR. Boolean networks are governed by a binary rule-based dynamic system, where nodes *x*_1_, *x*_2_, …, *x*_*n*_ take a value of 1 or 0 (ON/OFF), respectively. In the current implementation of BioSWITCH, we support Boolean logic because of its popularity in the context of GRNs due to the switch-like behaviour naturally exhibited by biological systems during the regulation of their functional states (Leshi et al., 2018).

State transitions, from 0 to 1, or from 1 to 0, are influenced by the node’s parent interactions, where relationships are either positive or negative. For instance, if *x*_*i*_ has *k*_*i*_ parent nodes 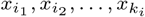, then the value of *x*_*i*_ at time *t* + 1 is determined by its parent nodes’ status at time *t*, described by the Boolean function:

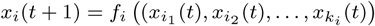

Assuming that the Boolean functions are time invariant, it follows that:

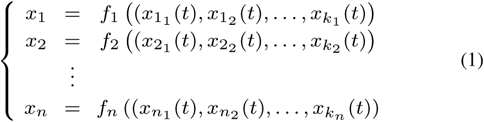

The global state of the network at any time point is the collection of states, as defined by 0 or 1, of each of the nodes. Thus, a network of *N* nodes, can undertake 2^*N*^ possible states, where the vector state is defined as:

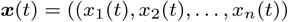

and the state space as:

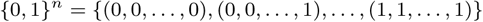

If we consider a network of three genes, *a, b, c*, its transitions would thus be defined by the functions:

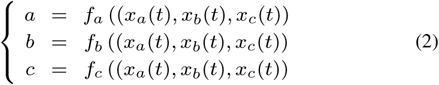

The 2^3^ possible states such network can undertake is schematically represented in Figure 1c, with stable states depicted in red. If reached, stable states are those where the dynamics of the network do not change over time. In the context of GRNs, the stable states translate biologically into the predicted cellular phenotype. BioSWITCH, via the GINML format, helps bridge a gap between BioPAX models and a simulation environments, such as GINSim, thereby enabling the calculation of such stable states, which allow us to predict the behaviour of network when different perturbation are applied *in silico*, such as the knock-out/inhibition (0) or activation (1) of a gene.

### 2.3 Merging and Expanding Gene Regulatory Networks

The increasing wealth and size of available network models calls for the need to automate the process of merging information from individual networks into unified, more comprehensive, networks. As such, we have provided the functionality to combine GINML networks into BioSWITCH, setting a starting point for the scalability of network models. Specifically, the *MergeTree* class compares the node names in a case-insensitive manner. If the nodes already exist, only the missing edges are added along with their corresponding rules, if the nodes do not exist in the original network, both nodes and edges are added to the graph along with any corresponding rules.

### 2.4 Applications

*In silico* methodologies are clearly becoming indispensable tools to navigate the ever growing volume of biological data. BioSWITCH enables the implementation of static BioPAX network data into a user-friendly executable simulation environment which is easily accessible to non-developers. The expansion of GINML-encoded networks and the addition of deposited knowledge to existing executable models has been done manually, BioSWITCH addressed the need to automate this process.

Importantly, we and others have recently developed models using the ABM framework, which can incorporate, amongst others, GRNs. Among these, *microC* models these networks into an ABM 3D spatial model of a cell, and execute them under different perturbation to infer cellular behaviour (Voukantsis et al., 2019). As input *microC* accepts the GXL-GINML format, which makes BioSWITCH-converted networks executable within cellular agents. BioSWITCH can be applied to either scale agents’ networks and/or integrate newly generated regulatory graphs from non-GINML databases in an increasingly automated fashion.

## 3 Results

To demonstrate and validate BioSWITCH functionality, we used a BioPAX-encoded GRN controlling haematopoiesis from Haydarlou et al. (2016), for which previously published simulation results were available. Most importantly, this model was validated by the BioPAX validator (Rodchenkov et al., 2013). Haematopoiesis, the process of blood formation, is an intensely studied process mainly controlled by transcriptional regulation. Bonzanni et al. (2013) created a Boolean network model, linking 11 transcription factors that govern this highly complex process. This original model was developed and simulated in Petri-net, another commonly used mathematical modelling language to describe discrete events of a dynamic system over time (Latorre-Biel and Jimenez-Macias, 2018). A BioPAX version of this model was later created by Haydarlou et al. (2016) for BioASF. We used this deposited version to test BioSWITCH faithfulness in reproducing the existing model. Namely, we converted the BioPAX deposited model into an executable GINML file. This proved that we were able to reconstruct the same network to be viewed and manipulated in GINSim (cf. Figure 1d). We then computed stable states, and could verify that they were identical to previously published simulation results in Haydarlou et al. (2016) and Bonzanni et al. (2013). Moreover, the newly generated haematopoiesis model resulted in the three following stable states: all 11 components set to 0, where GATA1 and Scl are the only expressed transcription factors, and a cyclic attractor of 32 substates. Finally, building on this example, we demonstrate how BioSWITCH enables us to combine the graph in Figure 1d with a smaller network of three nodes to output a larger network which can be seen in Figure 1e.

## 4 Discussion

Molecular systems orchestrating the biology of the cell involve a complex network of interactions among various components (i.e. genes, proteins, molecules). They can be studied *in silico* using a class of discrete models where the behaviour of the network translates into the long-term expected biological behaviour or cellular phenotype (Xiao, 2009). BioSWITCH provides access to a user-friendly interface for non-developers to run asynchronous and synchronous simulation experiments of BioPAX models. Specifically, it enables the simulation of BioPAX models within the GINSim environment. Using BioSWITCH we successfully converted an existing haematopoiesis BioPAX model to an executable Boolean logical GINML file. This allowed us to validate the BioSWITCH-generated model against previously validated published simulations of the same. We could verify that GINSim simulation of the GINML haematopoiesis model reproduced the same three stable states, including a cyclic attractor consisting of 32 substates, as the BioASF and Petri-Net simulation results. In addition, BioSWITCH contributes to the collection of GINML models to be reused by the scientific community and offers a starting point for the automatic expansion of existing GINSim network models.

This is not without limitations, however, since BioPAX files can contain various levels of information. BioSWITCH relies on generalised classes and rules which currently limits its use to BioPAX models with control interactions of type *TemplateReactionRegulation*, thus limiting the breadth of BioPAX models that can be tapped into. BioSWITCH served as a proof of concept and we now aim to expand this tool to other BioPAX models, newer releases will address these current limitations.

Importantly, we have already integrated BioSWITCH into an established ABM microC framework (Voukantsis et al., 2019), enabling us to integrate newly generated regulatory graphs into existing 3D cellular agents. Generating executable gene networks to build computational sub-agents will contribute to a deeper appreciation of the effect molecular changes have in a given environment, and may help elucidate mechanisms behind cell behaviour observed in different experimental models. BioSWITCH opens the door to the automatic generation of such computational representations by ‘switching on’ *static* network models thereby providing a new perspective from which to study their dynamics and response to perturbations *in silico*.

## Acknowledgements

We are grateful for the advice received from the BioPAX community and Pathway Commons founders, in particular Dr. Augustin Luna from the Dana-Farber Cancer Institute at Harvard Medical School. We also thank Dr. Charalampos Triantafyllidis for testing the BioSWITCH software.

## References

Voukantsis D, Kahn K, Hadley M, Wilson R, Buffa F. Modelling genotypes in their microenvironment to predict single- and multi-cellular behaviour. GigaScience 2019;giz010.

Letort G, Montagud A, Stoll G, Heiland R, Barillot E, Macklin P, et al. PhysiBoSS: a multi-scale agent-based modelling framework integrating physical dimension and cell signalling. Bioinformatics 2019;35(7):1188–1196.

Fabregat A, Sidiropoulos K, Garapati P, Gillespie M, Hausmann K, Haw R, et al. The Reactome pathway Knowledgebase. Nucleic Acids Res 2016;44:D481–D487.

Kutmon M, Riutta A, Nunes N, Hanspers K, Willighagen EL, Bohler A, et al. WikiPathways: capturing the full diversity of pathway knowledge. Nucleic Acids Research 2016;44:D488–D494.

Cerami EG, Gross BE, Demir E, Rodchenkov I, Babur O, Anwar N, et al. Pathway Commons, a web resource for biological pathway data. Nucleic Acids Res 2011;39:D6850–D690.

Kanehisa M, Furumichi M, Tanabe M, Sato Y, Morishima K. KEGG: new perspectives on genomes, pathways, diseases and drugs. Nucleic Acids Res 2017;45:D353–D361.

Demir E, Cary M, S P, et al. The BioPAX community standard for pathway data sharing. Nature Biotechnology 2010;28:935–942.

Horrocks I, Patel-Schneiderb PF, Harmelenc F. From SHIQ and RDF to OWL: the making of a Web Ontology Language. Web Semantics: Science, Services and Agents on the World Wide Web 2003;1(1):7–26.

Babur O, Dogrusoz U, Demir E, Sander C. ChiBE: interactive visualization and manipulation of BioPAX pathway models. Bioinformatics 2010;26(3):429–431.

Shannon P OOBNWJRDANSBIT Markiel A. Cytoscape: a software environment for integrated models of biomolecular interaction networks. Genome Research 2003;13(11):2498–2504.

Demir E, Babur O, Rodchenkov I, Aksoy BA, Fukuda KI, Gross B, et al. Using Biological Pathway Data with Paxtools. PLoS Comput Biol 2013;9.

Luna A, Babur O, Aksoy BA, Demir E, Sander C. PaxtoolsR: pathway analysis in R using Pathway Commons. Bioinformatics 2016;32(8):1262–1264.

Haydarlou R, Jacobsen A, Bonzanni N, Feenstra K, Abeln S, Heringa J. BioASF: a framework for automatically generating executable pathway models specified in BioPAX. Bioinformatics 2016;32:i60–i69.

Naldi A, Faure DBA, Lopez F, Thieffry D, Chaouiya C. Logical modelling of regulatory networks with GINsim 2.3. Biosystems 2009;97(2):134–139.

Holt RC, Schurr A, Sim SE, Winter A. GXL: A graph-based standard exchange format for reengineering. Science of Computer Programming 2006;60:149–170.

Leshi C, Don K, Sandhya S. A Novel Data-Driven Boolean Model for Genetic Regulatory Networks. Frontiers in Physiology 2018;9:1328.

Rodchenkov I, Demir E, Sander C, Bader GD. The BioPAX Validator. Bioinformatics 2013;29(20):2659–2660.

Bonzanni N, A G, Feenstra K, Schutte J, Kinston S, Miranda-Saavedra D, et al. Hard-wired heterogeneity in blood stem cells revealed using a dynamic regulatory network model. Bioinformatics 2013;29:i80–i88.

Latorre-Biel JI, Jimenez-Macias E. Petri Net Models Optimized for Simulation, Simulation Modelling Practice and Theory. IntechOpen 2018;.

Xiao Y. A tutorial on analysis and simulation of boolean gene regulatory network models. Current Genomics 2009;10:511–525.

